# Allosteric receptor modulation uncovers an FFAR2 antagonist as a positive orthosteric modulator/agonist in disguise

**DOI:** 10.1101/2021.05.19.444808

**Authors:** Simon Lind, Dagny Olofsson Hoffmann, Huamei Forsman, Claes Dahlgren

## Abstract

Two earlier described Free Fatty Acid Receptor 2 (FFAR2)-specific antagonists (CATPB and GLPG0974) have different receptor-interaction characteristics at the molecular/functional level. The inhibitory effect of the two antagonists, on the novel receptor-cross-talk activation signals generated by the ATP-receptor, show that both antagonists inhibit the effect of the positive allosteric FFAR2 modulators (PAMs) AZ1729 and Cmp58. No neutrophil activation was induced by AZ1729 or Cmp58 alone, but together they were co-agonistic PAMs and activated the superoxide generating NADPH-oxidase in neutrophils. This response was inhibited by CATPB but not by GLPG0974; in contrast, GLPG0974 acted as a positive modulator that increased the potency but not the efficacy of the response. At the signaling level, GLPG0974 changed the biased signaling induced by the co-agonistic PAMs, to include a rise in the cytosolic concentration of free calcium ions (Ca^2+^). This effect was reciprocal, i.e., GLPG0974 triggers a rise in intracellular Ca^2+^, demonstrating that GLPG0974 may act as an FFAR2 agonist. In summary, by studying the effects of the FFAR2 ligand GLPG0974 on neutrophils activation induced by the co-agonists AZ1729 and Cmp58, we reveal that GLPG0974 in addition to be an antagonist, displays also agonistic and positive FFAR2 modulating functions that affects the NADPH-oxidase activity and the receptor down-stream signaling induced by the two co-agonistic PAMs.

## 1. Introduction

Neutrophils are phagocytic white blood cells that are of prime importance in our innate immune system as well as in the initiation and resolution of aseptic inflammation [1]. The functions of these cells are regulated both by so called pathogen-associated molecular patterns (PAMPs) and damage-associated molecular patterns (DAMPs) originating from microbes and damaged host tissues, respectively. Many such inflammatory mediators are recognized by surface receptors that belong to the family of G protein-coupled receptors (GPCRs) [2, 3]. Members of this receptor family are all membrane proteins that change their basic conformation when agonists (activating ligands) bind to the so called orthosteric binding site exposed on the membrane surface of receptor-expressing cells [4, 5]. Free fatty acid receptor 2 (FFAR2 [6, 7]) is such a receptor expressed in neutrophils. This receptor recognizes short fatty acids (FFAs) such as acetate and propionate, metabolites produced by gut microorganisms during fermentation of dietary fibers. These metabolites are regarded not only as energy sources but also as regulators of inflammatory activities [8, 9]. The conformational changes induced by a receptor-bound agonist involves the receptor parts that are facing the cytoplasmic side of the membrane and initiate the down-stream signaling events. These receptor changes are in large prevented by conventional receptor specific antagonists, a group pharmacological tool-compounds of importance to dampen the responses induced by receptor agonists of endogenous or exogenous origin. Referring to FFAR2, two potent antagonists, GLPG0974 (4-[[(*R*)-1-(benzo[*b*]thiophene-3-carbonyl)-2-methyl-azetidine-2-carbonyl]-(3-chlorobenzyl)amino]-butyric acid); [10] and CATPB ((*S*)-3-(2-(3-chlorophenyl)acetamido)-4-(4-(trifluoromethyl)phenyl) butanoic acid; [11], have been shown to classify as conventional antagonists that compete with receptor specific agonists for binding to the orthosteric site, and these antagonists are highly selectivity for FFAR2 [12, 13].

The signals generated down-stream of the agonist-occupied receptor, initiated by the structural change of the receptor-domains on the cytosolic side of the receptor-expressing membrane, are determined by the properties of the activating agonist. Accordingly, the pharmacology and the structural changes adopted by a receptor is complex, a complexity that includes the variability and classification of agonists that interact with a receptor [14–16]. One type of agonists activates the receptor and amplify several different signaling pathways (the signaling and functional response is balanced), whereas the signaling and functional response induced by other agonists, that activate the same receptor, is biased (functionally selective); that is, one particular signaling pathway is preferred over another [15]. To add in complexity, receptor specific ligand may interact with a site on the receptor that is structurally and physically separated from the orthosteric binding pocket. Such ligands may modulate receptor functions positively or negatively, in that when the modulated receptor is activated by an orthosteric agonist, signaling of such receptors is not the same as that induced by the same agonist in the absence of the modulator [17]. An allosteric receptor modulator is expected to affect solely the activity induced by orthosteric agonists recognized by the allosterically modulated receptor [18, 19]. Although a receptor can harbor two or more allosteric binding sites specific for different allosteric modulators [20], these binding sites do not normally intercommunicate to activate the targeted receptor. GPCR-modulation/activation have, however, recently been shown to be more complex when it was shown that, i) allosteric modulators specific for FFAR2 affect signaling by this receptor but also by agonists specific for other neutrophil GPCRs such as the ATP-receptor P2Y_2_R and the formyl peptide receptors (FPRs) [21, 22], and that ii) the allosterically modulated FFAR2 may be transferred to an active biased signaling state by a second positive allosteric modulator (PAM) recognized by the same receptor. This activation is, thus, achieved without involvement of an orthosteric agonist [23, 24]. It is, thus, obvious that the down-stream signaling and receptor specific as well as non-specific receptor effects of allosteric receptor modulators are much more complex than initially anticipated.

It is clear from activation/inhibition patterns that characterize for example the opioid system, that ligands may affect receptor functions differently, as illustrated by the fact that a partial activating ligand may be a functional antagonist for activation with a full agonist. In addition, an activating ligand (agonist) for one receptor may be an inhibiting ligand (antagonist) for another receptor [25, 26]. Ligands possessing such a mixed activation(agonist)/inhibition(antagonist)-pattern have also been described for the formyl peptide receptor (FPRs) system. This is illustrated by data showing that agonists that specifically activate the murine version of Fpr2 may be antagonist for the human ortholog FPR2 [27, 28]. Accordingly, it is reasonable to assume that the structural change induced by an allosteric FFA2R modulator, that distinctly affect the affinity/efficacy of orthosteric agonist, also could affect the function of classical receptor antagonists that interact with the orthosteric binding site. The two receptor-specific FFAR2 antagonists mentioned (GLPG0974 and CATPB) have been shown to interact with the orthosteric binding site in FFAR2, and we have now investigated the effects of these antagonists on neutrophil FFAR2-activation when induced by two PAMs/co-agonists. The novel neutrophil activation mediated by the two allosteric FFA2R modulators/co-agonists, was largely reduced by CATPB but substantially augmented by GLPG0974. Taken together, we show that when FFAR2 is interdependently activated by two PAMs/co-agonists, GLPG0974 has no longer any antagonist effect but instead positively modulates the FFAR2 mediated response. In addition, the two interdependent PAMs turn GLPG0974 into an activating FFAR2 ligand (agonist) that induces a transient rise in the cytosolic concentration of free calcium ions.

## 2. Material and Methods

### 2.1. Chemicals

Isoluminol, TNF-α, ATP, propionic acid, bovine serum albumin (BSA), and fMLF were purchased from Sigma (Sigma Chemical Co., St. Louis, MO, USA). Cyclosporin H was a kind gift provided by Novartis Pharma (Basel, Switzerland). Dextran and Ficoll-Paque were obtained from GE-Healthcare Bio-Science (Uppsala, Sweden). Fura 2-AM was from Molecular Probes/Thermo Fisher Scientific (Waltham, MA, USA), and horseradish peroxidase (HRP) was obtained from Boehringer Mannheim (Mannheim, Germany). The allosteric FFAR2 modulator AZ1729 [29] was a generous gift from AstraZeneca (Mölndal, Sweden) and the phenylacetamide compound (S)-2-(4-chlorophenyl)-3,3-dimethyl-N-(5-phenylthiazol-2-yl)butanamide (PA;Cmp58 [30]) was obtained from Calbiochem-Merck Millipore (Billerica, USA). The FFAR2 antagonists GLPG0974 (4-[[(*R*)-1-(benzo[*b*]thiophene-3-carbonyl)-2-methyl-azetidine-2-carbonyl]-(3-chlorobenzyl)amino]-butyric acid) and CATPB ((S)-3-(2-(3-chlorophenyl)acetamido)-4-(4-(trifluoromethyl)-phenyl) butanoic acid), as well as the P2Y_2_R antagonist AR-C118925XX (5-[[5-(2,8-Dimethyl-5*H*-dibenzo[*a*,*d*]cyclohepten-5-yl)-3,4-dihydro-2-oxo-4-thioxo-1(2*H*)-pyrimidinyl]methyl]-*N*-2*H*-tetrazol-5-yl-2-furancarboxamide) were obtained from Tocris (Abingdon, UK). Subsequent dilutions of receptor ligand and other reagents were made in Krebs-Ringer Glucose phosphate buffer (KRG; 120 mM NaCl, 4.9 mM KCl, 1.7 mM KH_2_PO_4_, 8.3 mM NaH_2_PO_4_, 1.2 mM MgSO_4_, 10 mM glucose, and 1 mM CaCl2 in dH_2_O, pH 7.3).

### 2.2. Isolation of human neutrophils

Neutrophils were isolated from buffy coats from healthy blood donors by dextran sedimentation and Ficoll-Paque gradient centrifugation as described by Bøyum [31]. Remaining erythrocytes were removed by hypotonic lysis and the neutrophils were washed and resuspended in KRG. To amplify the activation signals the neutrophils were primed with TNF-α (10 ng/mL for 20 min at 37°C), and then stored on ice until use.

### 2.3. Measuring NADPH-oxidase activity

Isoluminol-enhanced chemiluminescence (CL) was used to measure superoxide, the precursor of production of other reactive oxygen species (ROS), generated by the neutrophil NADPH-oxidase [32, 33]. In short, the measurements were performed in a six-channel Biolumat LB 9505 (Berthold Co., Wildbad, Germany), using disposable 4-ml polypropylene tubes and a 900 μl reaction mixture containing 10^5^ neutrophils, isoluminol (2×10^−5^ M) and HRP (4 Units/ml). The tubes were equilibrated for 5 min at 37°C, before addition of agonist (100 μl) and the light emission was recorded continuously over time. In experiments where the effects of receptor specific antagonists were determined, these were added to the reaction mixture 1-5 min before stimulation. Control neutrophils incubated under the same condition but in the absence of an antagonist were run in parallel for comparison.

### 2.4. Calcium mobilization

Neutrophils at a density of 1–3×10^6^ cells/ml were washed with Ca^2+^-free KRG and centrifuged at 220x*g*. The cell pellets were re-suspended at a density of 2×10^7^ cells/ml in KRG containing 0.1% BSA, and loaded with 2 μM FURA 2-AM for 30 min at room temperature. The cells were then washed once with KRG and resuspended in the same buffer at a density of 2×10^7^cells/ml. Calcium measurements were carried out in a Perkin Elmer fluorescence spectrophotometer (LC50), with excitation wavelengths of 340 nm and 380 nm, an emission wavelength of 509 nm, and slit widths of 5 nm and 10 nm, respectively. The transient rise in intracellular calcium ([Ca^2+^]_i_) is presented as the ratio of fluorescence intensities (340 nm / 380 nm) detected.

### 2.5 Statistical analysis

Statistical calculations were performed in GraphPad Prism 8.02 (Graphpad Software, San Diego, CA, USA). The specific statistical tests are stated in the relevant figure legend. A *p*-value < 0.05 was regarded as statistically significant and is indicated by \**p* < 0.05, \*\**p* < 0.01. Statistical analysis was performed on raw data values using a one-way ANOVA followed by Dunnett’s multiple comparison or, paired Student’s *t*-test.

### 2.6. Ethics Statement

In this study, conducted at the Sahlgrenska Academy in Sweden, buffy coats obtained from the blood bank at Sahlgrenska University Hospital, Gothenburg, Sweden have been used. According to the Swedish legislation section code 4§ 3p SFS 2003:460 (Lag om etikprövning av forskning som avser människor), no ethical approval was needed since the buffy coats were provided anonymously and could not be traced back to a specific donor.

## 3. Results

### 3.1. FFAR2 antagonists inhibit the effects of receptor specific modulators in neutrophils activated without any orthosteric FFAR2 agonist

The non-activating allosteric FFAR2 modulators Cmp58 and AZ1729, turn the orthostertic agonist propionate into an efficient activator of the neutrophil superoxide (O_2_^−^) generating NADPH-oxidase (Fig 1A). In addition, the non-activating P2Y_2_ receptor (P2Y_2_R) specific agonist ATP as well as non-activating concentrations of the FPR1 specific agonist fMLF were turned to activating agonists in neutrophils with FFAR2 allosterically modulated (Fig 1B and E). The response induced by ATP/fMLF required the presence of an allosteric FFAR2 modulator and was achieved through a receptor cross-talk activation mechanism without the involvement of any orthosteric FFAR2 agonist (for details regarding the receptor cross-talk activation mechanism see [3, 21]). These activation systems were used to further investigate the effects of FFAR2 specific antagonists on the interaction of the PAMs Cmp58 and AZ1729 with FFAR2.

**Figure 1.**
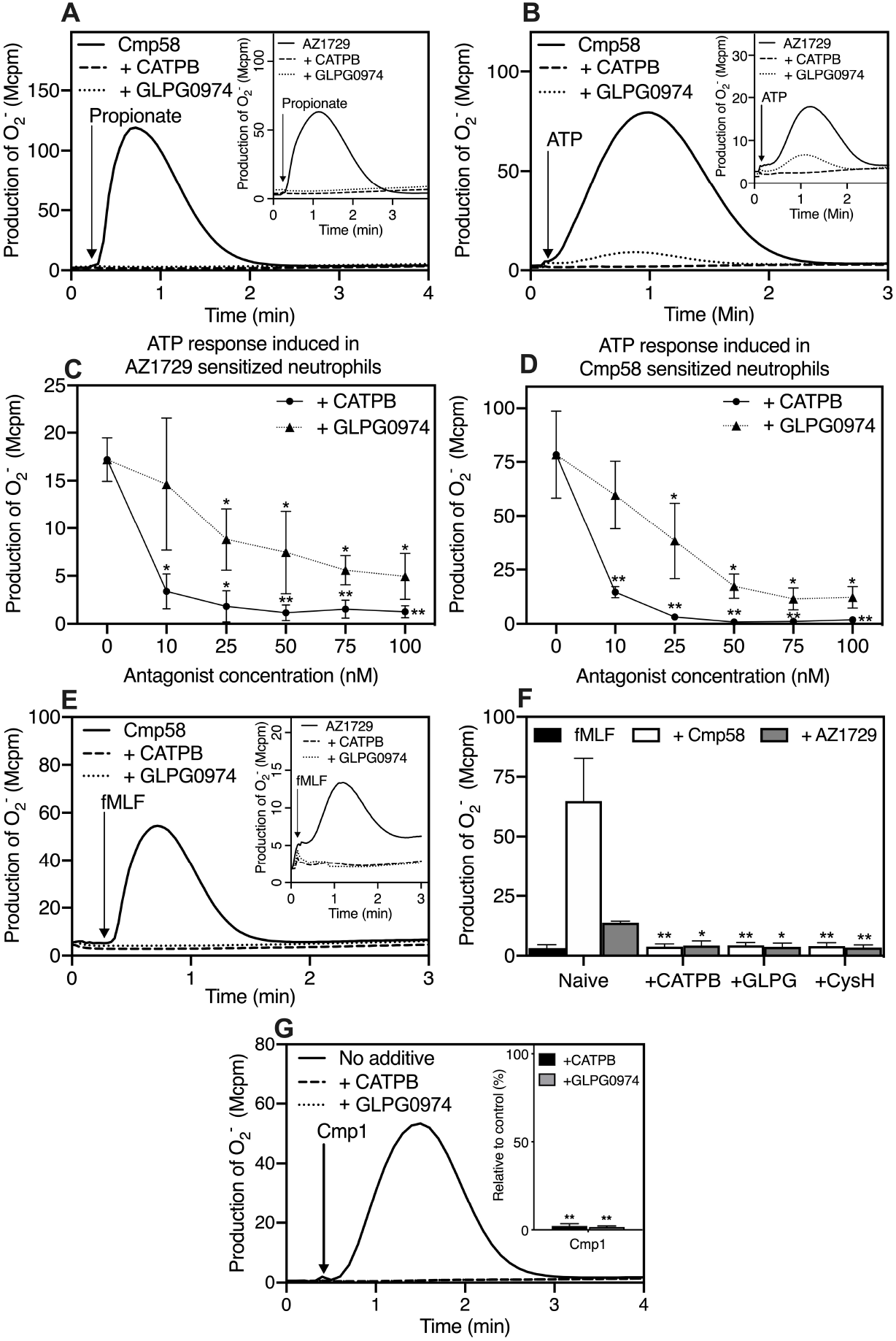
Activation of the neutrophil superoxide (O_2_^−^) generating NADPH-oxidase and effects of the allosteric FFAR2 modulators Cmp58 and AZ1729. **(A)** Production of O_2_^−^ in neutrophils sensitized either with Cmp58 (1 μM for 5 min; solid line) or AZ1729 (1 μM for 5 min; solid line; inset) when activated by propionate (25 μM; time point for addition indicated by an arrow) in the absence or presence of an FFAR2 specific antagonist (CATPB, dashed line, or GLPG0974, dotted line; 100 nM). One representative experiment out of > 5 is shown. **(B)** Production of O_2_^−^ in neutrophils sensitized either with Cmp58 (1 μM for 5 min; solid line) or AZ1729 (1 μM for 5 min; solid line; inset) when activated by ATP (10 μM; time point for addition indicated by an arrow) in the absence or presence of an FFAR2 specific antagonist (CATPB, dashed line, or GLPG0974, dotted line; 100 nM). One representative experiment out of > 5 is shown. **(C)** Production of O_2_^−^ in neutrophils sensitized with Cmp58 (1 μM for 5 min; **C**) or AZ1729 (1 μM for 5 min; **D**) when activated by ATP (10 μM) in the absence or presence of different concentrations of an FFAR2 specific antagonist, CATPB (solid line) or GLPG0974 (dotted line). The responses were determined from the peak activities and expressed in Mcpm (mean ± SD, n = 3). Statistical analyses were performed using a one-way ANOVA followed by a Dunnett’s multiple comparison test comparing the peak responses in the absence and presence of respective inhibitor. **(E)** Production of O_2_^−^ in neutrophils sensitized either with Cmp58 (1 μM for 5 min; solid line) or AZ1729 (1 μM for 5 min; solid line; inset) when activated by the FPR1 specific agonist fMLF (1 nM) in the absence (solid line) or presence of an FFAR2 specific antagonist, CATPB, (dashed line; 100 nM) or GLPG0974 (dotted line; 100 nM). One representative experiment out of > 5 is shown and time for addition of the agonist is marked by an arrow. **(F)** Production of O_2_^−^ in neutrophils sensitized either with Cmp58 (1 μM for 5 min) or AZ1729 (1 μM for 5 min) when activated by the FPR1 specific agonis fMLF (1 nM) in the absence or presence of an FFAR2 specific antagonist (CATPB or GLPG0974; 100 nM) and an FPR1 specific antagonist (cyclosporin H; 1μM), respectively. The response induced by fMLF in the absence of any modulator is shown for comparison (black bar to the left). The responses were determined from the peak activities and expressed in Mcpm; mean ± SD, n = 3. Statistical analyses were performed using a one-way ANOVA followed by a Dunnett’s multiple comparison test comparing the peak responses in the absence and presence of respective inhibitor. **(G)** Production of O_2_^−^ in neutrophils when activated by the FFAR2 specific agonist Cmp1 (1 μM; time point for addition indicated by an arrow) in the absence (solid line) or presence of an FFAR2 specific antagonist, CATPB (dashed line, 100 nM) or GLPG0974 (dotted line, 100 nM). One representative experiment out of > 5 is shown and time for addition of the agonist is marked by an arrow. **Inset:** Effects of specific FFAR2 antagonists (CATPB and GLPG0974; 100 nM) on neutrophil production of O_2_^−^ induced by the FFAR2 agonist Cmp1. The responses were determined from the peak activities and expressed in percent of the remaining activity in the presence of an antagonist (mean ± SD, n = 3). Statistical analyses were performed using a one-way ANOVA followed by a Dunnett’s multiple comparison test comparing the peak responses in the absence and presence of respective inhibitor.

As expected, and in agreement with previous findings, the response induced by propionate, in neutrophils allosterically modulated by Cmp58 (Fig 1A) and AZ1729 (Fig 1A inset), was fully inhibited by CATPB and GLPG0974, two well-known specific FFAR2 antagonists. The direct neutrophil activation (without any PAM) induced by Cmp1, a potent small compound FFAR2 agonist [11] was also fully inhibited by the two FFAR2 antagonists (Fig 1G). Moreover, also the response induced by ATP was inhibited by either of the two antagonists (Fig 1B). We show that the two FFAR2 specific antagonists CATPB and GLPG0974, earlier described to be recognized by the same receptor site as conventional orthosteric FFAR2 agonists, inhibit the effect of the allosteric modulators AZ1729 and Cmp58 when the activation signals were triggered by ATP. A potent inhibition was obtained with a concentration ratio between FFAR2 modulator and antagonist of around 20 for both Cmp58/CATPB and AZ1729/CATPB (Fig 1C and D). Despite the fact that both antagonists reduced the ATP induced response, no full inhibition was reached using GLPG0974 at a concentration ratio of modulator/antagonist of 10, with either of the two allosteric modulators. The response was reduced by around 65 and 85 percent with the allosteric modulator AZ1729 and Cmp58, respectively, and this degree of inhibition was reached with the highest concentration of GLPG0974 used (Fig 1C and D). The two FFAR2 antagonists CATPB and GLPG0974 also inhibited the allosteric FFAR2 modulators when the activation signals were triggered by fMLF (Fig 1E and F). The two FFAR2 antagonists, thus, inhibit not only neutrophil activation triggered by the orthosteric agonists propionate and Cmp1, but also the response induced by ATP/fMLF, responses that require the presence of an allosteric FFAR2 modulator and is achieved without any orthosteric agonist. Thus, the FFAR2 antagonists directly inhibit the effects of the allosteric modulators Cmp58 and AZ1729.

### 3.2. CATPB inhibits whereas GLPG0974 positively modulates the response induced by the allosteric modulators/co-agonists AZ1729 and Cmp58

The structural change needed to activate the signaling pathways down-stream of an allosterically modulated receptor could in principle be achieved not only by an orthosteric agonist, but also by another allosteric modulator recognized by a second allosteric binding site exposed by the same receptor. Accordingly, we have shown that the two non-activating FFAR2 PAMs Cmp58 and AZ1729 co-operate to activate neutrophils [23, 24]. When added together the two modulators potently activated the superoxide producing NADPH-oxidase (Fig 2). Despite the fact that the two FFAR2 antagonists potently reduced the activity of the Cmp58- and AZ1729-dependent responses (see Fig 1), a very small inhibition was obtained with CATPB (Fig 2A) and basically no inhibition was obtained with GLPG0974 (Fig 2B), in neutrophils activated by the combined effect of the two co-agonistic PAMs (1 μM each; Fig 2A and B).

**Figure 2.**
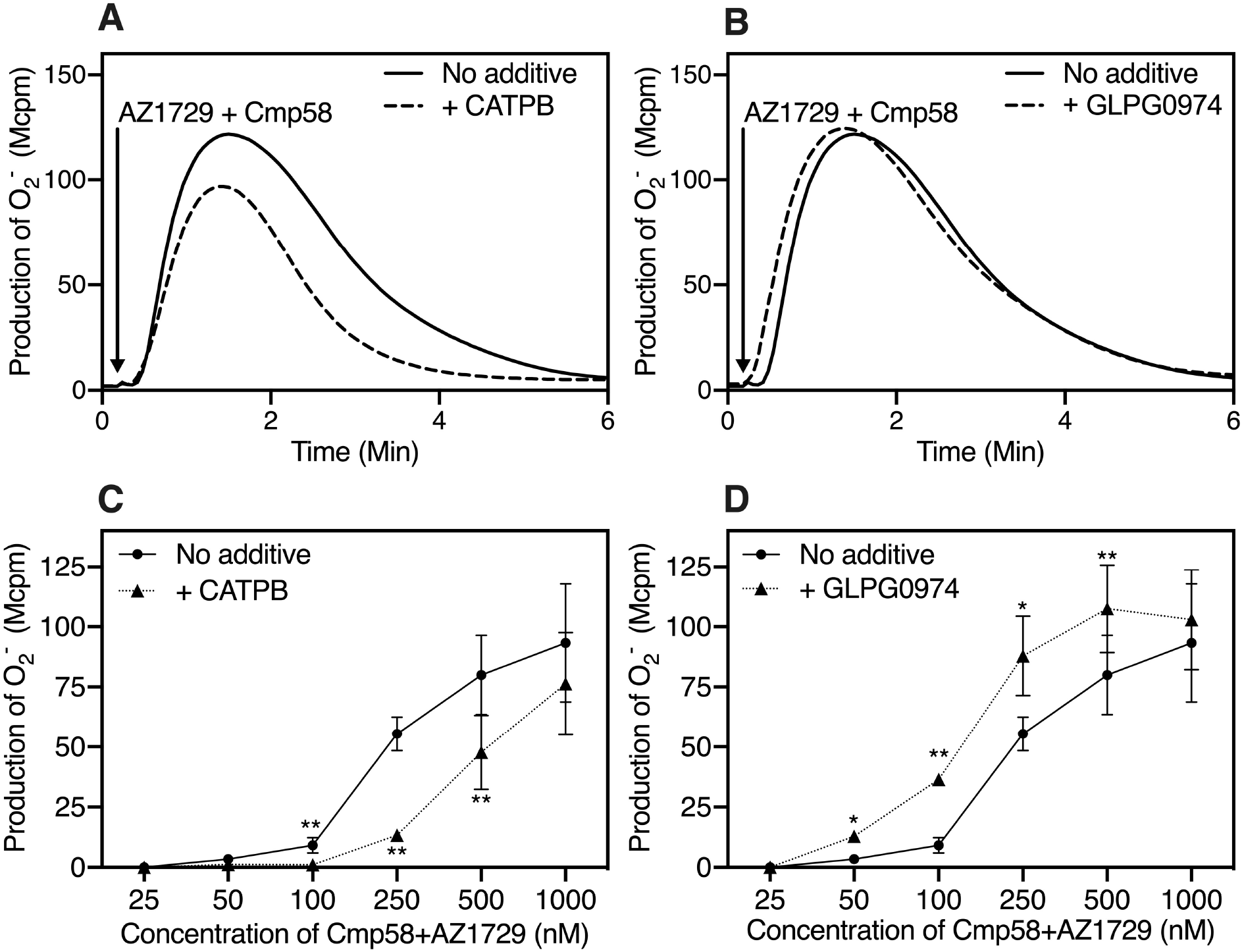
Activation of the neutrophil superoxide (O_2_^−^) generating NADPH-oxidase when induced together by the allosteric FFAR2 modulators AZ1729 and Cmp58 and effects of the FFAR2 antagonists CATPB and GLPG0974. **(A)** Production of O_2_^−^ in neutrophils activated with the co-agonists Cmp58 (1 μM) and AZ1729 (1 μM) in the absence (solid line) or presence of the FFAR2 specific antagonist CATPB (dashed line, 100 nM). One representative experiment out of > 5 is shown and time for addition of the co-agonists is marked by an arrow. **(B)** Production of O_2_^−^ in neutrophils activated with the co-agonists Cmp58 (1 μM) and AZ1729 (1 μM) in the absence (solid line) or presence of the FFAR2 specific antagonist GLPG0974 (dashed, 100 nM). One representative experiment out of > 5 is shown and time for addition of the co-agonists is marked by an arrow. **(C)** Production of O_2_^−^ in neutrophils activated with different concentrations of the co-agonists Cmp58 (25 nM-1 μM) and AZ1729 (25 nM-1 μM) in the absence (solid line) or presence of the FFAR2 specific antagonist CATPB (dotted line, 100 nM). The responses were determined from the peak activities and expressed in Mcpm (mean ± SD, n = 3). The statistical analysis was performed using paired Student’s *t*-test comparing the peak responses induced either by absence or presence of the antagonist CATPB when activating with different concentrations of the co-agonists Cmp58 and AZ1729. **(D)** Production of O_2_^−^ in neutrophils activated with different concentrations of the co-agonists Cmp58 (25 nM-1 μM) and AZ1729 (25 nM-1 μM) in the absence (solid line) or presence of the FFAR2 specific antagonist GLPG0974 (dotted line, 100 nM). The responses were determined from the peak activities and expressed in Mcpm (mean ± SD, n = 3). The statistical analysis was performed using paired Student’s *t*-test comparing the peak responses induced either by absence or presence of CATPB when activating with different concentrations of the co-agonists Cmp58 and AZ1729.

The amount of O_2_^−^ produced was dependent on the concentration of the two co-agonistic modulators with an EC_50_ value around 200 nM each, and it is clear that the effects of the antagonists changed when the concentrations of the activating allosteric modulators were reduced. In agreement with the expected effect of an antagonist, CATPB inhibited the response induced by Cmp58/AZ1729 when the concentrations of the two co-agonists were reduced. A good inhibition was obtained at a concentration of the co-agonistic modulators of 250 nM each (Fig 2C). No corresponding effect on the response was, however, obtained with GLPG0974 (Fig 2D) when the concentrations of the co-agonists were reduced. On the contrary, the response was increased in the presence of this antagonist (Fig 2D). The positive modulating effect of GLPG0974 was evident at a 500 nM concentration of the two co-agonists and the effect was most pronounced at a concentration of 100 nM each of Cmp58/AZ1929, reaching levels five times (or more) higher than the naive (non-amplified) response. The co-agonists AZ1729 and Cmp58, that interdependently activate the neutrophil NADPH-oxidase, thus, transfer the FFAR2 specific ligand GLPG0974 from a receptor antagonist to a positive FFAR2 modulator that affects the potency (i.e., the EC_50_ value) but not the efficacy (i.e., the E_max_ value) of the response (Fig 2D).

### 3.3. The positive modulating effect of GLPG0974 is more pronounced when concentration of either of the allosteric modulators AZ1729 and Cmp58 was reduced

Keeping the concentration of AZ1729 constant (1 μM), the amount of O_2_^−^ produced was dependent on the concentration of the other co-agonist/modulator with an EC_50_ value of around 20 nM for Cmp58 (Fig 3A). As shown above, no effect of GLPG0974 was obtained when the neutrophil response was induced by Cmp58/AZ1729 at a concentration of 1 μM each, and no inhibition of the response obtained when the concentration of Cmp58 was reduced to 100 nM (Fig 3A). The AZ1729/Cmp58 induced response was, however, increased by GLPG0974 at lower concentrations of Cmp58. The positive modulating effect of GLPG0974 was most pronounced at really low concentrations of Cmp58 (Fig 3A).

**Figure 3.**
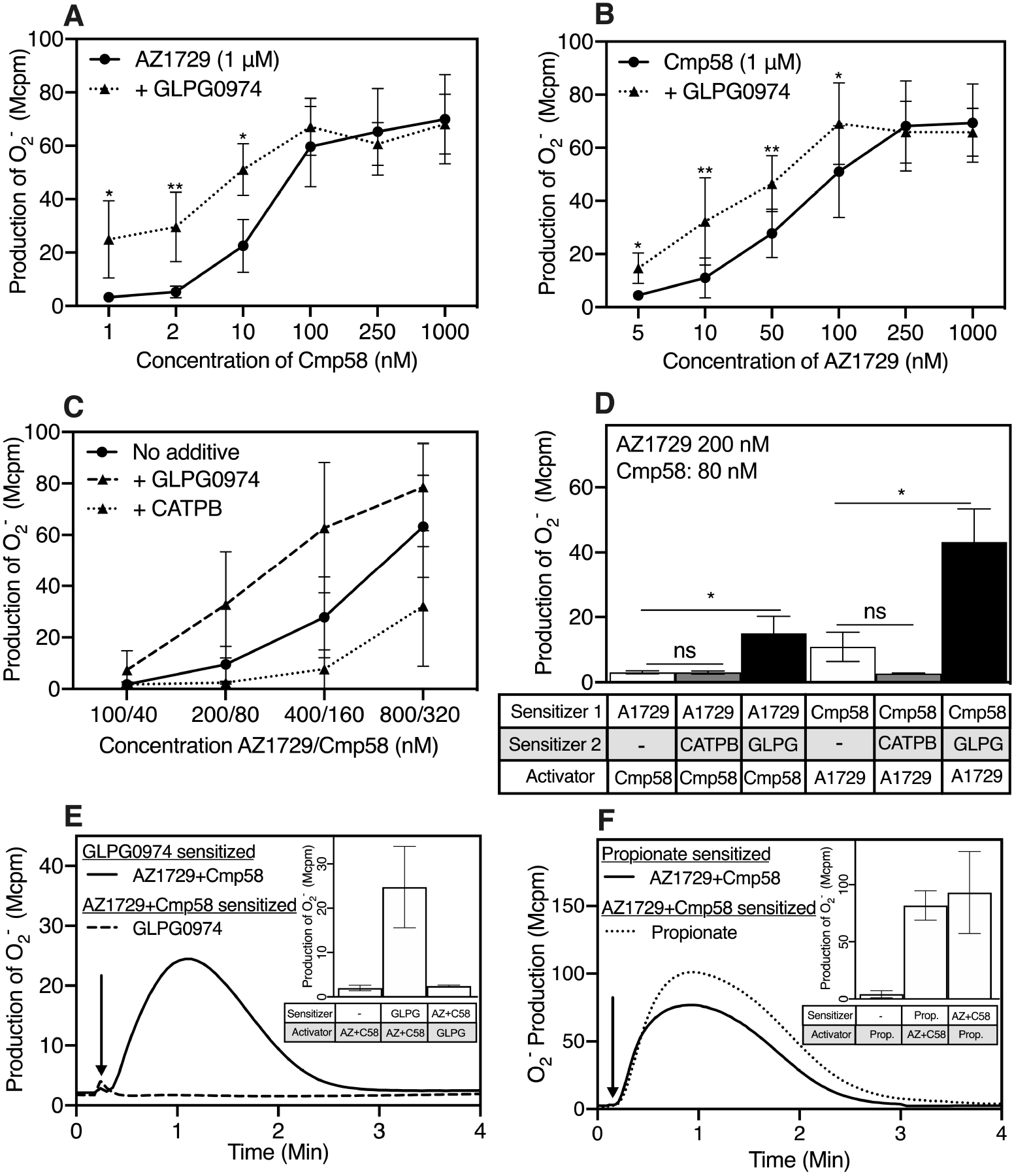
Activation and inhibition patterns in neutrophils activated by the allosteric FFAR2 modulators/co-agonists Cmp58 and AZ1729. **(A)** Production of O_2_^−^ in neutrophils activated with a fixed concentration of the co-agonist AZ1729 (1 μM) and different concentration of Cmp58 (1 nM-1 μM) in the absence (solid line) or presence of the FFAR2 specific antagonist GLPG0974 (dotted line, 100 nM). The responses were determined from the peak activities and expressed in Mcpm; mean ± SD, n = 3. The statistical analysis was performed using paired Student’s *t*-test comparing the peak responses induced either by absence or presence of the antagonist GLPG0974. **(B)** Production of O_2_^−^ in neutrophils activated with a fixed concentration of the co-agonist Cmp58(1 μM) together with different concentration of AZ1729 (1 nM-1 μM) in the absence (solid line) or presence of the FFAR2 specific antagonist GLPG0974 (dotted line, 100 nM). The responses were determined from the peak activities and expressed in Mcpm; mean ± SD, n = 3. The statistical analysis was performed using paired Student’s *t*-test comparing the peak responses induced either by absence or presence of the antagonist GLPG0974. **(C)** Production of O_2_^−^ in neutrophils activated with variable concentrations of the co-agonists Cmp58 (40-320 nM) together with variable concentration of AZ1729 (100-800 nM) in the absence (solid line) or presence of the FFAR2 specific antagonists GLPG0974 (broken line, 100 nM) or CATPB (dotted line, 100 nM). The responses were determined from the peak activities and expressed in Mcpm; mean ± SD, n = 3. **(D)** Neutrophils were sensitized with AZ1729 (200 nM) or Cmp58 (80 nM) alone (White bars) or combined with either CATPB (grey bars, 100 nM) or GLPG0974 (black bars, 100 nM). The AZ1729 sensitized neutrophils were activated by Cmp58 (80 nM) whereas the Cpm58 sensitized were activated by AZ1729 (200 nM) and the production of O_2_^−^ was measured. The responses were determined from the peak activities and expressed in Mcpm; mean ± SD, n = 3. Statistical analyses were performed using a one-way ANOVA followed by a Dunnett’s multiple comparison test comparing the peak responses in the absence and presence of either CATPB or GLPG0974. **(E)** Neutrophils were sensitized with non/low activating concentrations of AZ1729/Cmp58 (5 min, 200 and 80 nM, respectively) or with GLPG0974 (5 min, 100 nM). The GLPG0974 sensitized neutrophils were activated with AZ1729/Cmp58 (200 and 80 nM respectively, solid line) whereas the AZ1729/Cmp58 sensitized neutrophils were activated by GLPG0974 (100 nM, broken line) and the production of O_2_^−^ was measured. One representative experiment out of > 5 is shown and time for addition of the co-agonists/GLPG0974 is marked by an arrow. **Inset:** Neutrophils were sensitized with AZ1729 (200 nM) together with Cmp58 (80 nM) or with GLPG0974 (100 nM). The AZ1729/Cmp58 sensitized neutrophils were activated by GLPG0974 (100 nM) and the GLPG0974 sensitized were activated by AZ1729/Cmp58 (200 nM/80 nM), respectively, and the production of O_2_^−^ was measured. For control, non-sensitized neutrophils were also activated by the co-agonist AZ1729 (200 nM) combined with Cmp58 (80 nM). The responses were determined from the peak activities and expressed in Mcpm (mean ± SD, n = 3). **(F)** Neutrophils were sensitized with non/low activating concentrations of AZ1729/Cmp58 (5 min, 200 and 80 nM, respectively) or with propionate (5 min, 25 μM). The propionate sensitized neutrophils were activated with AZ1729/Cmp58 (200 and 80 nM respectively, solid line) whereas the AZ1729/Cmp58 sensitized neutrophils were activated by propionate (25 μM, broken line) and the production of O_2_^−^ was measured. One representative experiment out of > 5 is shown and time for addition of the co-agonists/GLPG0974 is marked by an arrow. **Inset:** Neutrophils were sensitized with AZ1729 (200 nM) combined with Cmp58 (80 nM) or propionate (25 μM). The AZ1729/Cmp58 sensitized neutrophils were activated by propionate (25 μM) and the propionate sensitized were activated by AZ1729/Cmp58 (200 nM/80 nM), respectively, and the production of O_2_^−^ was measured. For control, non-sensitized neutrophils were also activated by propionate alone (25 μM). The responses were determined from the peak activities and expressed in Mcpm (mean ± SD, n = 3).

In agreement with the data presented above, the amount of O_2_^−^ produced at a constant concentration of Cmp58 (1 μM) was also dependent on the concentration of AZ1729 with an EC_50_ value for this allosteric modulator of around 50 nM (Fig 3B). No effect was obtained with GLPG0974 on the response obtained when the concentration of AZ1729 was lowered to 250 nM (Fig 3B). The AZ1729/Cmp58 induced response was, however, substantially increased by GLPG0974 at concentrations of AZ1729 lower that 250 nM. The positive modulating effect of GLPG0974 was most pronounced at even lower concentrations of AZ1729 (Fig 3B).

It is obvious from the data presented that in comparison to AZ1729, lower concentration of Cmp58 can be used to activate the neutrophil NADPH-oxidase, but the results were the same when different concentrations of the two co-agonists were used to activate neutrophils, that is, the inhibitory effect of CATPB increased when the concentrations of the co-agonists/modulators were reduced whereas the relative potentiating effect of GLPG0974 increased when the concentrations of the activating ligands were reduced (Fig 3C).

### 3.4. The positive modulating effect of GLPG0974 is not reciprocal

The allosteric modulator Cmp58 activates neutrophils sensitized with AZ1729, and this response is positively modulated by GLPG0974. The same response pattern was obtained with AZ1729 when used to activate neutrophils sensitized with Cmp58 (Fig 3D). It should also be noticed that neutrophils sensitized with GLPG0974 were activated by the co-agonistic modulators AZ1729 and Cmp58 in concentrations so low that they were non-activating in the absence of GLPG0974 (Fig 3E); no activation was, however, induced by GLPG0974, when added to neutrophils sensitized with the non-activating concentrations of the modulators (Fig 3E inset). Propionate could replace GLPG0974 as the sensitizing compound (Fig 3F), but with propionate, this activation was reciprocal. Accordingly, propionate was turned to activating agonist in neutrophils sensitized with the two modulators, but in addition, non-activating concentrations of the co-agonistic modulators potently activated neutrophils sensitized with propionate (Fig 3F).

### 3.5. GLPG0974 expands signaling down-stream of FFAR2 in neutrophils activated by two co-agonistic allosteric receptor modulators

High concentrations of propionate activate the PLC-PIP_2_-IP_3_-Ca^2+^-signaling pathway in neutrophils, and in agreement with the inhibition characteristics of the two orthosteric FFAR2 antagonists CATPB and GLPG0974, this transient rise in [Ca^2+^]_i_ was inhibited by both antagonists (Fig 4A). Despite the activation of the neutrophil NADPH-oxidase induced by the two allosteric FFA2R modulators (co-agonists) AZ1729 and Cmp58 when added together, this was not accompanied by any change in the free concentration of [Ca^2+^]_i_) (Fig 4A, inset). The inability to trigger a rise in [Ca^2+^]_i_ was evident, both when the two modulators were added together, and when added in sequence. The biased signaling pattern induced by the co-operative action of the two allosteric FFA2R modulators/co-agonists, characterized by an activation of the NADPH-oxidase without any activation of the PLC-PIP_2_-IP_3_-Ca^2+^- signaling pathway was, however, changed by GLPG0974. GLPG0974 was without effect on the activation of the neutrophil NADPH-oxidase induced by high concentrations of AZ1727/Cmp58, but a transient rise in [Ca^2+^]_i_ was induced by two co-agonists in neutrophils sensitized with GLPG0974 (Fig 4B). No such effect was obtained with CATPB (Fig 4B).

**Fig 4.**
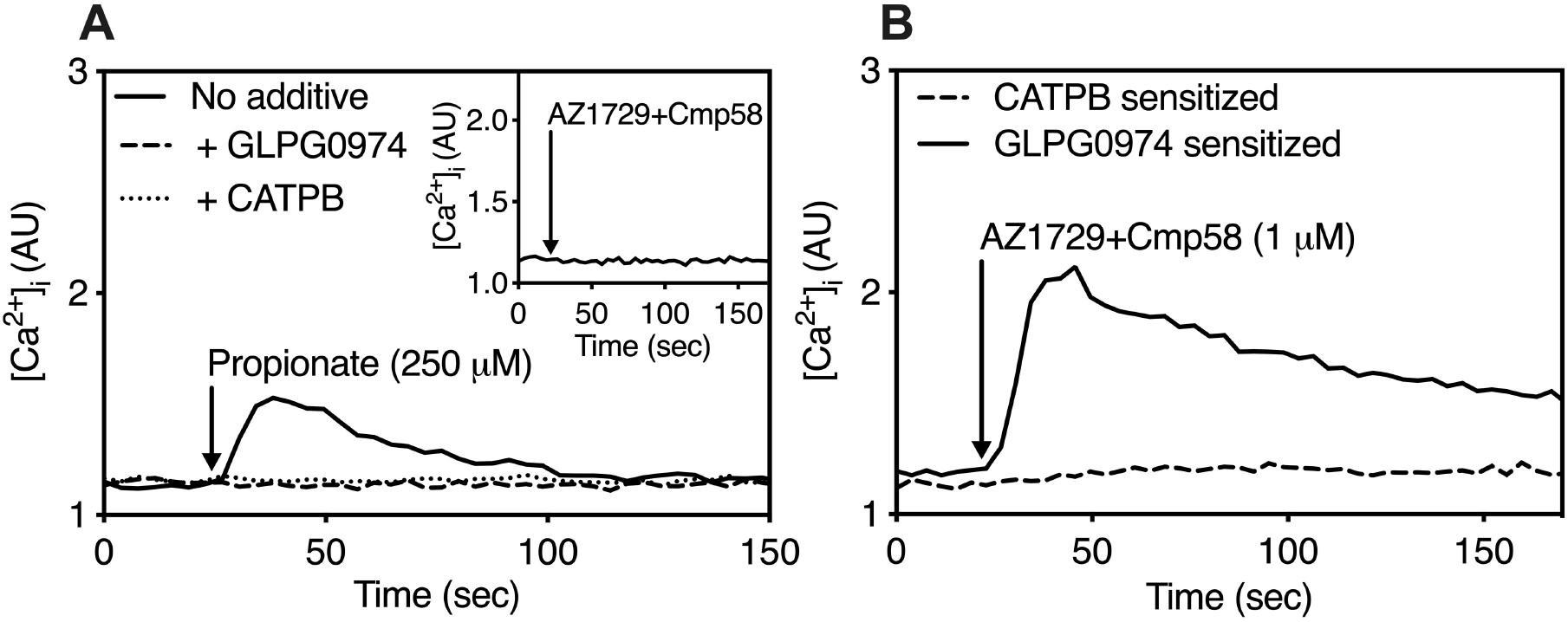
Effects of FFAR2 antagonists on the rise in intracellular concentration of Ca^2+^([Ca^2+^]_i_) in neutrophils. **(A)** The antagonist-mediated inhibition of the change in [Ca^2+^]_i_ was followed in neutrophils activated by propionate (250 μM). Neutrophils were preincubated without or with an antagonist, CATPB (dotted line, 100 nM) or GLPG0974 (dashed line, 100 nM) and the cells were then activated by propionate. The time point for addition of propionate is indicated by an arrow. One representative experiment out of 3 independent experiment is shown. Abscissa, time of study (sec) Ordinate, the increase in [Ca^2+^]_i_ is expressed as the ratio between Fura-2 fluorescence at 340 nm and 380 nm. Inset: No transient rise in [Ca^2+^]_i_ was induced by the co-agonistic allosteric modulators Cmp58 (1 μM) and AZ1729 (1 μM). The time point for addition of Cmp58/AZ1729 is indicated by an arrow. **(B)** The change in [Ca^2+^]_i_ induced by AZ1729 (1 μM) combined with Cmp58 (1 μM) was followed in neutrophils sensitized with the FFAR2 antagonists CATPB (dashed, 100 nM) and GLPG0974 (dotted line, 100 nM), respectively. The time points for addition of AZ1729/Cmp58 (1 μM each) is indicated by arrows. One representative experiment out of 3 independent experiment is shown. Abscissa, time of study (sec) Ordinate, the increase in [Ca^2+^]_i_ is expressed as the ratio between Fura-2 fluorescence at 340 nm and 380 nm.

No transient rise in [Ca^2+^]_i_ was induced in GLPG0974 sensitized neutrophils by either of the PAMs alone (Fig 5A). Both modulators are, thus, needed for the response. This is also illustrated by the fact that a rise in [Ca^2+^]_i_ was obtained in GLPG0974 sensitized neutrophils also when the two modulators were added in sequence (Fig 5A). Taken together, we show that the Cmp58/AZ1729-induced signaling is transferred from being biased to be balanced; the signals that activate NADPH-oxidase were in GLPG0974 sensitized neutrophils coupled also to a rise [Ca^2+^]_i_.

**Figure 5.**
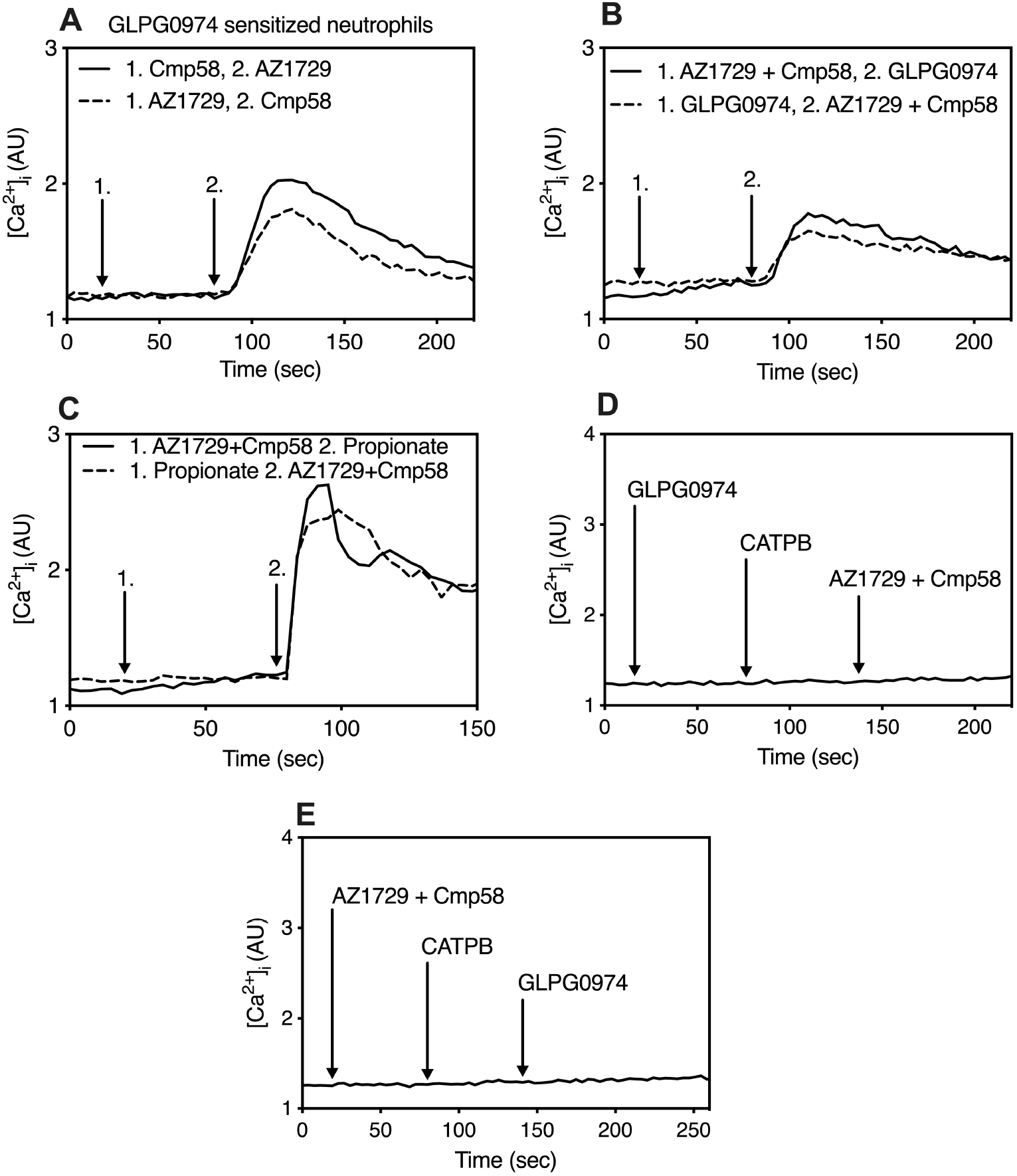
Response patterns following activation by the two allosteric modulators AZ1729 and Cmp58. **(A)** The change in [Ca^2+^]_i_ induced by AZ1729 (1 μM) and Cmp58 (1 μM) added in sequence to GLPG0974 (100 nM)-sensitized neutrophils. The time points for addition of the allosteric modulators are indicated by arrows. One representative experiment out of 3 independent experiment is shown. Ordinate, the increase in [Ca^2+^]_i_ is expressed as the ratio between Fura-2 fluorescence at 340 nm and 380 nm. **(B)** The transient rise in [Ca^2+^]_i_ induced by i) GLPG0974 (100 nM) in neutrophils sensitized with AZ1729/Cmp58 (1 μM each) and ii) by AZ1729/Cmp58 (1 μM each) in neutrophils sensitized with GLPG0974 (100 nM). The time points for addition of the different compounds are indicated by arrows. One representative experiment out of 3 independent experiment is shown. Ordinate, the increase in [Ca^2+^]_i_ is expressed as the ratio between Fura-2 fluorescence at 340 nm and 380 nm. **(C)** The transient rise in [Ca^2+^]_i_ induced by i) propionate (25 μM) in neutrophils sensitized with AZ1729/Cmp58 (1 μM each) and ii) AZ1729/Cmp58 (1 μM each) in neutrophils sensitized with propionate (25 μM). The time points for addition of the different compounds are indicated by arrows. One representative experiment out of 3 independent experiment is shown. Ordinate, the increase in [Ca^2+^]_i_ is expressed as the ratio between Fura-2 fluorescence at 340 nm and 380 nm. **(D)** Inhibition by CATPB (100 nM) of the transient rise in [Ca^2+^]_i_ induced by AZ1729/Cmp58 (1 μM each) in GLPG0974 (100 nM) sensitized neutrophils. The time points for addition of the different compounds are indicated by arrows. **(E)** Inhibition by CATPB (100 nM) of the transient rise in [Ca^2+^]_i_ induced by GLPG0974 (100 nM) in AZ1729/Cmp58 (1 μM each) sensitized neutrophils. The time points for addition of the different compounds are indicated by arrows. One representative experiment out of 3 independent experiment is shown. Ordinate, the increase in [Ca^2+^]_i_ is expressed as the ratio between Fura-2 fluorescence at 340 nm and 380 nm.

### 3.6. The FFAR2 antagonist GLPG0974 is an agonist in neutrophils sensitized/activated by the two co-agonistic allosteric FFAR2 modulators AZ1729 and Cmp58

Cmp58/AZ1729-induced signaling is, thus, transferred by GLPG0974 to be balanced and include a rise [Ca^2+^]_i_. This response was also reciprocal, as GLPG0974 triggered a rise in [Ca^2+^]_i_ in neutrophils sensitized with AZ1729/Cmp58 (Fig 5B). The allosteric modulators, unable to trigger a rise in [Ca^2+^]_i_ in neutrophils, lowered the thresh-hold for the propionate response, and also this response was reciprocal (Fig 5C). The response induced by Cmp58/AZ1729 in neutrophils sensitized with GLPG0974 was inhibited by CATPB (Fig 5D) and the same effect of this antagonist was obtained when the order of the sensitizing and activating ligands was reversed (Fig 5E).

The data show that the antagonist GLPG0974 is transferred to a ligand that activates the Ca^2+^-signaling rout in AZ1729/Cmp58 sensitized neutrophils, and this response was inhibited by the other FFAR2 antagonist CATPB. Since the NADPH-oxidase activity induced by the co-operative action of AZ1729 and Cmp58 was very pronounced, two experimental set-ups with activating and non-activating concentrations, respectively, of the PAMs were compared to determine the differences/similarities between GLPG0974 and propionate to also activate the neutrophil NADPH-oxidase. In neutrophils sensitized with low/non-activating concentrations of Cmp58/AZ1729, no activation was induced by GLPG0974 whereas propionate was a potent activator of the oxidase (see Fig 3 E and F).

At higher concentrations of Cmp58/AZ1729, the NADPH-oxidase activity induced peaked after a fairly short time period and the response was then rapidly terminated. Following termination of the response, the neutrophils were homologously desensitized as no response was induced when a new dose of AZ1729/Cmp58 was added (data not shown). No activation of the oxidase was induced by GLPG0974 (Fig 6A) or propionate (Fig 6B) when added to neutrophils once the AZ1729/Cmp58 induced response was back to background levels. Taken together these data suggest that the PAMs Cmp58/AZ1729 transfer GLPG0974 to a functional selective FFAR2 activating ligand that favors the Ca^2+^-signaling route over the NADPH-oxidase activating pathway.

**Figure 6.**
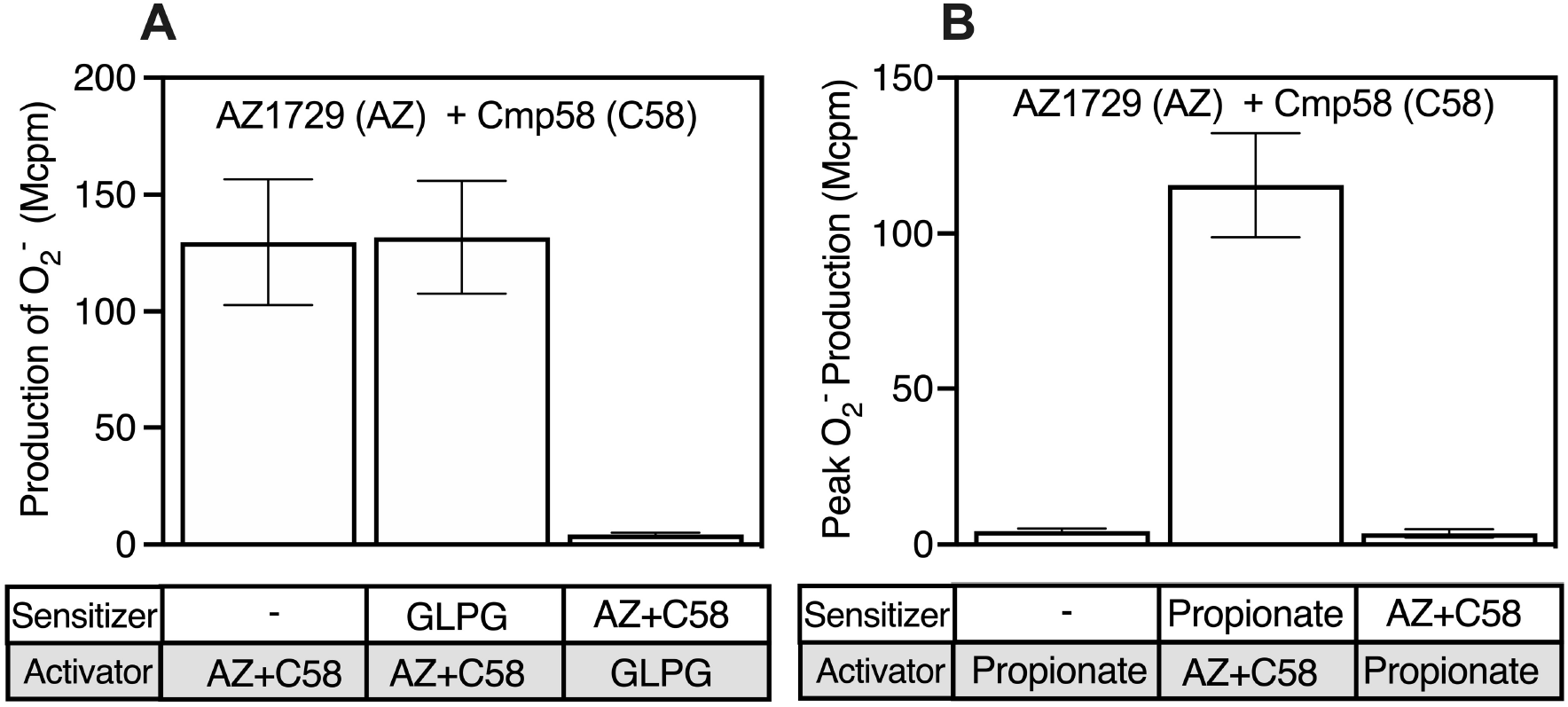
Desensitization of the GLPG0974 and the propionate induced responses. **(A)** Production of O_2_^−^ in neutrophils sensitized either with AZ1729/Cmp58 (1 μM each) or GLPG0974 (100 nM). The AZ1729 sensitized neutrophils were activated with GLPG0974 (100 nM) and the GLPG0974 sensitized neutrophils were activated with AZ1729/Cmp58 (1 μM each), respectively. For comparison, non-sensitized neutrophils were also activated with the co-agonist AZ1729/Cmp58 (1 μM each). The responses were determined from the peak activities and expressed in mega counts per minute (Mcpm) (mean ± SD, n = 3). **(B)** Production of O_2_^−^ in neutrophils sensitized either with AZ1729/Cmp58 (1 μM each) or propionate (25 μM). The AZ1729 sensitized neutrophils were activated with propionate (25 μM) and the propionate sensitized neutrophils were activated with AZ1729/Cmp58 (1 μM each), respectively. For comparison, non-sensitized neutrophils were also activated with propionate (25 μM each) and AZ1729/Cmp58 (1 μM each), respectively. Neutrophils was also stimulated with propionate (25 μM) alone. The responses were determined from the peak activities and expressed in mega counts per minute (Mcpm) (mean ± SD, n = 3).

### 3.7. No increase in [Ca^2+^]_i_ is induced by GLPG0974 in propionate desensitized neutrophils

Cmp58/AZ1729 lowered the thresh-hold for the propionate response and as expected, this response was inhibited by CATPB. In addition, no propionate response was induced in Cmp58/AZ1729 sensitized neutrophils first activated with GLPG0974 (Fig 7A). The same response pattern was obtained when the order by which the activating agonists added was changed. That is, no GLPG0974 response is induced in Cmp58/AZ1729 sensitized neutrophils first activated with propionate (Fig 7B). These results, showing that propionate inhibits the response induced by GLPG0974, suggest that the orthosteric FFAR2 agonist desensitizes the receptor for activation with GLPG0974.

**Figure 7.**
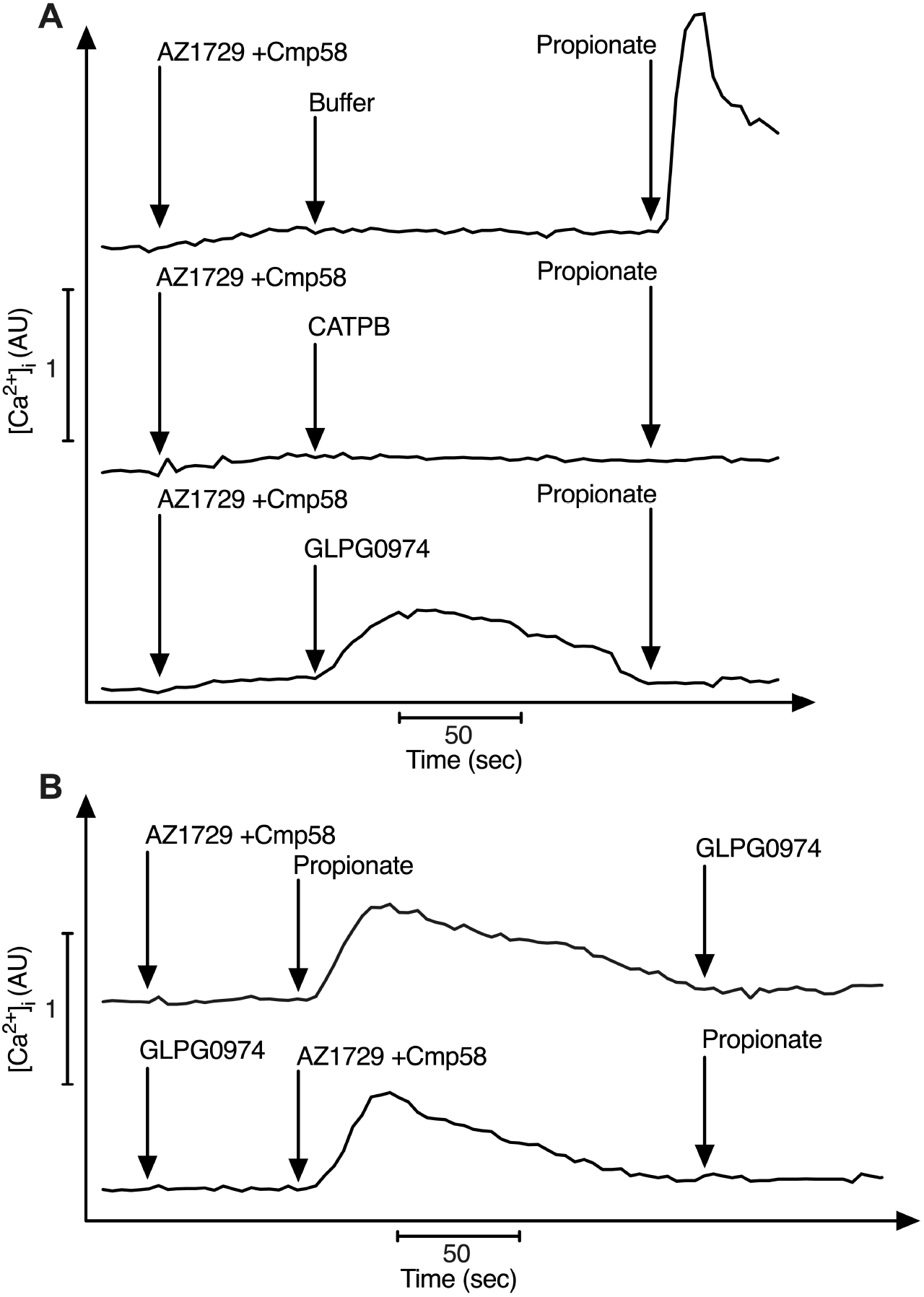
Desensitization of the GLPG0974 induced response. **(A)** The propionate induced change in [Ca^2+^]_i_ in AZ1729/Cmp58 (250 nM each) sensitized neutrophils (upper curve) was inhibited by CATPB (100 nM; middle curve) as well as in neutrophils first activated with GLPG0974 (100 nM; lower curve).The time points for addition of the different compounds are indicated by arrows. **(B)** No change in [Ca^2+^]_i_ is induced by GLPG0974 in AZ1729/Cmp58 (250 nM each) sensitized neutrophils first activated with propionate (25 μM; upper curve) and likewise, no change in [Ca^2+^]_i_ is induced by propionate (25 μM; lower curve) in GLPG0974 sensitized (100 nM) neutrophils first activated AZ1729/Cmp58 (250 nM each). The time points for addition of the different compounds are indicated by arrows. One representative experiment out of 3 is shown. Abscissa, time of study (the bar represents 50 s); Ordinate, the increase in [Ca^2+^]_i_ given as the ratio between Fura-2 fluorescence at 340 nm and 380 nm.

## 4. Discussion

A positive allosteric modulator (PAM) that interacts with a G protein-coupled receptor (GPCRs) promote conformational changes in the receptor, and by definition this interaction leads to changed signaling and functional responses induced by an orthosteric agonist that binds to the modulated receptor [34]. The PAM-induced changes have alone normally no direct signaling effects downstream of the modulated receptor, but the outcome is an increase in binding affinity for an agonist and/or of the maximum response (efficacy) induced by orthosteric agonists. It is reasonable to assume that PAMs, that by binding to an allosteric site affect the structure/function of the orthosteric binding site, also could affect the function of a receptor-specific antagonist that binds to the orthosteric receptor site. In agreement with this assumption, we show that GLPG0974, a free fatty acid receptor 2 (FFAR2) specific antagonist that binds to the orthosteric binding site of this receptor [12, 13], is turned into a mixed receptor modulator/agonist in neutrophil phagocytes activated by two earlier described co-agonistic PAMs, AZ1729 and Cmp58 [3, 23, 24].

The interaction and binding properties of the two FFAR2 specific antagonist (CATPB and GLPG0974) included in the study have previously been fairly well characterized/documented [10, 11, 13, 35, 36]. Although their detailed binding characteristics differ, it has been shown that both antagonists fit into, and bind specifically to the orthosteric site/binding pocket in human FFAR2 and that their respective affinity for the receptor are very similar or even identical [7, 13]. Accordingly, both antagonists inhibit the response induced by natural (i.e., short chain fatty acids) as well as synthetic (i.e., small compounds such as Cmp1; [11, 37] orthosteric FFAR2 agonists. The two PAMs AZ1729 and Cmp58 were originally identified in screening studies [29, 30] designed to identify specific FFAR2 ligands, and they both lack the carboxylic acid part suggested to be required for orthosteric FFAR2 agonists [13]. This implies that both are ligands that interact with allosteric receptor sites, which is in agreement with the fact that AZ1729 and Cmp58 alone have no direct neutrophil activating effects and the two exert very similar modulating functions with respect to their effects on the response induced by the orthostatic FFAR2 agonist propionate [23, 24]. However, in order to determine the direct inhibitory effects of the antagonists on the allosteric FFAR2 modulators, measuring systems have to be used, in which a FFAR2 mediated response was achieved without involvement of any orthosteric FFAR2 agonist. We have earlier shown that the two allosteric modulators in addition to the effects on the response induced by orthosteric agonists, also positively increase the neutrophil response to the P2Y_2_R agonist ATP as well as to agonists recognized by FPRs [21, 22]. Accordingly, this activation of FFAR2, achieved through a novel receptor cross-talk mechanism that activates FFAR2 without involvement of any orthosteric FFAR2 agonist [3], was used to define the inhibitory properties of the antagonists CATPB and GLPG0974. The two antagonists inhibit the activation signals triggered by ATP in the presence of either of the allosteric FFAR2 modulators AZ1729 or Cmp58. No full inhibition was, however, obtained with GLPG0974 but still, the antagonist inhibited the allosteric modulating effects of both AZ1729 (≈60% inhibition) and Cmp58 (≈80% inhibition). Taken together, the data presented suggests that the respective FFAR2 binding site for Cmp58 and AZ1729 partly overlap with the orthosteric binding site that recognizes also the FFAR2 specific antagonists. In accordance with this, the antagonist CATPB and GLPG0974 block the effects of both AZ1729 and Cmp58.

The two allosteric FFAR2 modulators Cmp58 and AZ1729, recognized by two distinct binding sites on the receptor, interdependently activate neutrophils (this study and [23, 24]. The down-stream signaling profile of FFA2R, when activated by the two allosteric modulators, is biased. No transient rise in [Ca^2+^]_i_ is triggered during this activation (this study and [23, 24]). The precise receptor sites involved in the modulation are not known, except that previous findings show that the two modulators interact with different allosteric binding sites overlapping the orthosteric site. In agreement with this, the FFA2R antagonist CATPB also inhibits the activation induced when the two allosteric modulators are combined. This response was, however, not inhibited by GLPG0974; in contrast, GLPG0974 amplified the response. The co-agonists AZ1729 and Cmp58, thus, transfer the FFAR2 specific ligand GLPG0974 from a receptor antagonist to a positive orthosteric FFAR2 modulator that affects the potency (i.e., the EC_50_ value) rather that the efficacy (i.e., the E_max_ value) of the response. FFAR2 signaling down-stream of FFAR2 was biased when activated by the co-agonists, in that no activation of the PLC-PIP_2_-IP_3_-Ca^2+^pathway was obtained. GLPG0974 changed this signaling profile. The data presented show, that in addition to the positively modulated effect mediated by the antagonist GLPG0974 on the oxidase activity triggered by the co-agonists AZ1729 and Cmp58, the initial biased receptor down-stream signaling induced by the two allosteric modulators was changed to also include a transient rise in [Ca^2+^]_i_. It should also be noticed that this activation of the Ca^2+^ pathway was reciprocal, meaning that GLPG0974 was turned into an activating agonist in neutrophils sensitized with Cmp58/AZ1729. The GLPG0974 induced response was inhibited by the CATPB but also by propionate, suggesting that the response is inhibited not only by an FFAR2 antagonist but also when the receptor is desensitized following activation with an orthosteric agonist.

Allosteric interactions should be reciprocal in nature and the results presented regarding Ca^2+^ signaling induced by AZ1729/Cmp58 and GLPG0974 are in agreement with the reciprocity characterizing the effects of allosteric modulators [19, 38]. Despite the fact that the interaction with the receptor differs between the two orthosteric ligands GLPG0974 and propionate, the same type of Ca^2+^ signaling reciprocity was obtained with the agonist propionate and the allosteric modulators AZ1729/Cmp58. The FFAR2 binding difference between the two orthosteric ligands has been clearly shown using a DREADD (Designer Receptor Activated only by Designer Drug)-mutant of FFAR2 [39]; GLPG0974 still binds the mutated receptor with high affinity and receptor signaling is inhibited, whereas basically no activation is achieved with propionate. In agreement with this, the signaling selectivity of GLPG0974 partly differs from that induced by the orthosteric FFAR2 agonist propionate. GLPG0974 signaling was selective, in that no activation of the NADPH-oxidase was induced by this ligand in AZ1729/Cmp58 sensitized cells, irrespectively of the concentrations of the PAMs/co-agonists. Similar to GLPG0974, no or a very low activation of the NADPH-oxidase was induced by propionate in neutrophils first co-operatively activated with AZ1729/Cmp58 (1 μM each). Propionate induced, however, a response in neutrophils sensitized with low/non-activating concentrations of AZ1729/Cmp58, and this activation was reciprocal. Irrespective of the precise modulation mechanism for the two allosteric modulators AZ1729 and Cmp58 (that is a higher affinity for the second ligand or lowered energy-barrier for the conformational shift to a signaling state), and for GLPG0974, the latter not only positive modulated the oxidase activity induced by the two PAMs; the signaling pattern down-stream of the activated FFAR2 was changed to include also a transient rise in [Ca^2+^]_i_.

Receptor agonists and antagonist are generally defined by their respective capacity activate a receptor and to inhibit binding/function of agonists recognized by the same receptor. The outcome of ligand interaction may, however, be more complex. Receptor ligands may have both functions and by that be classified as mixed agonists/antagonists. Such ligands are generally agonists that only partly activate its receptor, and when combined with an agonist that alone has a very strong (full) effect on the receptor activity, the antagonistic (inhibiting) effect of a mixed agonist/antagonist is disclosed [25, 26]. The data showing that the response patterns in the presence of an FFAR2 antagonist differ depending both on the antagonist and on the mechanism by which the receptor is activated, adds to the complexity of how to define a receptor specific ligand. One of the FFAR2 antagonists studied, CATPB, inhibited activation of the receptor and this effect was evident both when activation involved an orthosteric agonist and when it was achieved independent of such a ligand. Based on the large similarities, regarding structural and receptor interaction characteristics, between CATPB and GLPG0974, we expected similar or even identical inhibitory properties of the two antagonists. Accordingly, and as expected, also GLPG0974 inhibited activation when an orthosteric agonist was part of the process and when the allosterically modulated FFAR2 was activated by the receptor cross-talk signals generated by P2Y_2_R or FPR1. The activation of the superoxide generating neutrophil NADPH-oxidase, induced by the interdependent synergistic PAMs AZ1729 and Cmp58 was, however, positively modulated by GLPG0974. Even though it has been shown that CATPB and GLPG0974 are able to interact with the same arginine residues (Arg_180_ and Arg_255_) in the orthosteric binding pocket in FFAR2 and have binding modes and affinities that are fairly similar [12, 13, 39], their respective mode of interaction obviously differs. It remains to be determined, if the difference between the antagonists could be explained at the molecular level, by the fact that that the length of the carboxylate chain differ between the antagonists; GLPG0974 has a longer and more flexible carboxylate chain allowing this ligand to interact electrostatically with the presumably more flexible Arg_180_, whereas the shorter chain in CATPB facilitates an electrostatic interaction with Arg_255_ [12, 13]. The antagonist GLPG0974 is tuned not only into a positive orthosteric modulator in neutrophils activated by the allosteric modulators/co-agonists AZ1729 and Cmp58, but these FFAR2 ligand also turn the antagonist into a biased FFAR2 agonist that activates the PLC-PIP_2_-IP_3_-Ca^2+^signaling pathway but not the pathway leading to an activation of the NADPH-oxidase. It should be noticed that GLPG0974 has entered a clinical phase 2 trial for treatment of intestinal bowel disease but it was in that study shown to be non-efficacious [36]. The novel findings regarding the changed function of GLPG0974 by FFAR2 modulators adds to the complex pharmacology of this receptor (possibly also in different disease settings), and should opens for new interest in this receptor as a target in inflammatory diseases.

## Abbreviations

AZ1729 and Cmp58: allosteric FFAR2 modulators
BSA: bovine serum albumin
CATPB: an FFA2R antagonist
CL: chemiluminescence
FFAR2: free fatty acid receptor 2
GPCR: G protein-coupled receptor
HRP: horse radish peroxidase
ns: no significant difference
P2Y_2_R: receptor for ATP
KRG: Krebs-Ringer Glucose phosphate buffer
ROS: reactive oxygen species
TNF-α: tumor necrosis factor α

## Authorship contributions

*Planning and formulation of research goals and study design*: Lind, Dahlgren

*Conducted experiments and validations:* Lind, Olofsson Hoffman

*Data presentation and analysis:* Lind, Olofsson Hoffman, Forsman, Dahlgren

*Supervised the research:* Lind, Forsman

*Writing of original draft*: Lind, Dahlgren

*Contributed to the writing of the manuscript:* Lind, Olofsson Hoffman, Forsman, Dahlgren

## 5. Acknowledgements

The work was supported by the Swedish Medical Research Council (HF, 02448), the Clas Groschinsky Foundation (HF, MI562), the Swedish Foundation for Strategic Research (HF, SM-17-0046, Åke Wibergs Foundation (HF, M15-005), and the Swedish state under the ALF-agreement (CD, ALFGBG 72510; HF, 78150). The research funders did not have any role in any part of the study.

We thank the members of the Phagocyte Research Group at the Sahlgrenska Academy, University of Gothenburg, for critically discussing the results and the manuscript.

